# Optimized cryo-EM data acquisition workflow by sample thickness determination

**DOI:** 10.1101/2020.12.01.392100

**Authors:** Jan Rheinberger, Gert Oostergetel, Guenter P Resch, Cristina Paulino

## Abstract

Sample thickness is a known key parameter in cryo-electron microscopy (cryo-EM) and can affect the amount of high-resolution information retained in the image. Yet, common data acquisition approaches in single particle cryo-EM do not take it into account. Here, we demonstrate how the sample thickness can be determined before data acquisition, allowing to identify optimal regions and restrict automated data collection to images with preserved high-resolution details. This quality over quantity approach, almost entirely eliminates the time- and storage-consuming collection of suboptimal images, which are discarded after a recorded session or during early image processing due to lack of high-resolution information. It maximizes data collection efficiency and lowers the electron microscopy time required per dataset. This strategy is especially useful, if the speed of data collection is restricted by the microscope hardware and software, or if microscope access time, data transfer, data storage and computational power are a bottleneck.

**Synopsis:** Sample thickness is a key parameter in single particle cryo-electron microscopy. Determining sample thickness before data acquisition allows to target optimal areas and maximize data output quality of single particle cryo-electron microscopy sessions. Scripts and optimized workflows for EPU and SerialEM are presented and available as open-source.

## Introduction

In single particle cryo-electron microscopy (cryo-EM), the thin layer of vitreous ice embedding the protein or macromolecule complex of interest is a key parameter in sample preparation and optimization. Obtained by rapidly plunge-freezing grids with holey support films in liquid coolant (Adrian *et al*., 1984), it preserves the structural integrity of the macromolecular complex of interest. The electron transparency of the vitreous ice layer surrounding the specimen in transmission electron microscopy (TEM), critically depends on the layer thickness: the thicker the vitreous layer the more scattering events (elastic and inelastic) occur, up to a point where electrons can no longer penetrate. In addition, thicker layers experience a defocus gradient, which leads to a stronger dampening of higher frequencies, limiting the resolution of the final reconstruction obtained (Wu *et al*., 2016). Conversely, a too thin ice layer might not suffice to fully embed the protein, leading to denaturation at the air-water interface, or push the sample towards the edge of the support film hole, limiting the number of copies in the field of view (Noble *et al*., 2018; D’Imprima *et al*., 2019). The importance and urgency for the optimization of cryo-EM sample preparation is reflected in the development of new techniques and devices in recent years (Dandey *et al*., 2018; Rubinstein *et al*., 2019; Ravelli *et al*., 2020; Tan & Rubinstein, 2020; Arnold *et al*., 2017; Kontziampasis *et al*., 2019; Mäeots *et al*., 2020).

Determining the thickness of a specimen is nothing new and has been described previously (Suloway et al., 2005; Cho *et al*., 2013; Yan *et al*., 2015; Rice *et al*., 2018). Yet, it has been mostly calculated on high-resolution images and used to monitor sample thickness during an ongoing data collection, or to select micrographs based on their sample thickness after data acquisition. In contrast, our work focuses on using sample thickness determination at low magnification to set-up an automated thickness-based hole-targeting, restricting data collection to optimal regions, similar to what has been available in Leginon (Suloway et al., 2005). We have successfully integrated this approach, available as open-source, in the commonly used data acquisition software packages SerialEM (Mastronarde, 2005; Schorb *et al*., 2019) and EPU (Thermo Fischer Scientific), and have tested it on 200 and 300 kV high-end electron microscopes. During the revision of this manuscript, an update on new features for the data acquisition software Leginon describes a similar approach (Cheng *et al*., 2021), highlighting its importance and making this workflow now available for the vast majority of used software packages.

As previously described and implemented in Leginon (regions, similar to what has been available in Leginon (Suloway et al., 2005; Rice *et al*., 2018) two fundamentally different approaches to estimate the thickness from projection images exist, which depend on the configuration of the microscope. In the case of a microscope equipped with an energy filter, one can benefit from the thickness dependent number of inelastic scattering events. Due to the energy loss, these electrons will be removed by the filter when operated in the zero-loss mode.

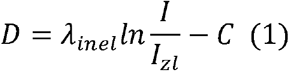

The thickness *D* is proportional to the *In* of the ratio between the mean intensity without (*I*) and with slit inserted (*I_zl_*) (Eq 1). The apparent mean-free-path λ_inel_ acts as scaling factor and describes the average distance electrons travel through the sample before an inelastic scattering event occurs. The correction term C, not present in the original description (Rice *et al*., 2018), is used by the scripts presented in this work to set the measurements in a hole over vacuum to zero.

For the aperture limited scattering (ALS) method (Eq. 2), sample thickness can be determined by comparing the mean intensity over vacuum (*I_0_*) versus the mean intensity over ice (*I*), utilizing the effect that the objective aperture removes part of the elastically scattered electrons. The number of scattered electrons is again proportional to the thickness (and content of the sample) (Angert *et al*., 1996; Cho *et al*., 2013; Rice *et al*., 2018). Here, instead of the mean-free-path, a scaling factor termed the ALS coefficient λ_ALS_ is used, which equally dependents on the acceleration voltage, objective aperture size, sample content (Rice *et al*., 2018) and imaging mode (LM vs M/SA).

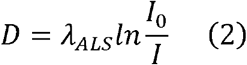

Both, λ_inel_ and λ_ALS_, can be determined experimentally by tomography or comparison between both methods (Noble *et al*., 2018; Rice *et al*., 2018).

In this work, we present our approach for an optimized data collection workflow by targeting only holes with optimal sample thickness. Thereby, we substantially increase data collection efficiency by maximizing output quality, while minimizing the amount of images collected that do not contribute to high-resolution information and which are discarded right away or early on during image processing. This is achieved by either combining Digital Micrograph (Gatan) scripts with the *Filter Ice Quality* histogram implemented in EPU (Thermo Fisher Scientific) for targeting, or by using scripts that combine the entire procedure in SerialEM (Mastronarde, 2005; Schorb *et al*., 2019). Based on quality assessments, such as resolution estimation of CTF fit (better than 4 Å), more than 90-95% of the data collected with this workflow provides high-resolution information.

## Results and discussion

### Digital Micrograph & EPU

With the functions available through Digital Micrograph (version 2.2 or higher), we developed a script that allows the user to calculate and monitor the sample thickness of a grid square with a holey-carbon support film at low magnification. The respective holes are colored with a heatmap as too-thick, optimal, or too-thin, based on a user-defined thickness range (Fig. 1 (a)). In addition, the user can determine the thickness at any point in the image.

**Figure 1.**
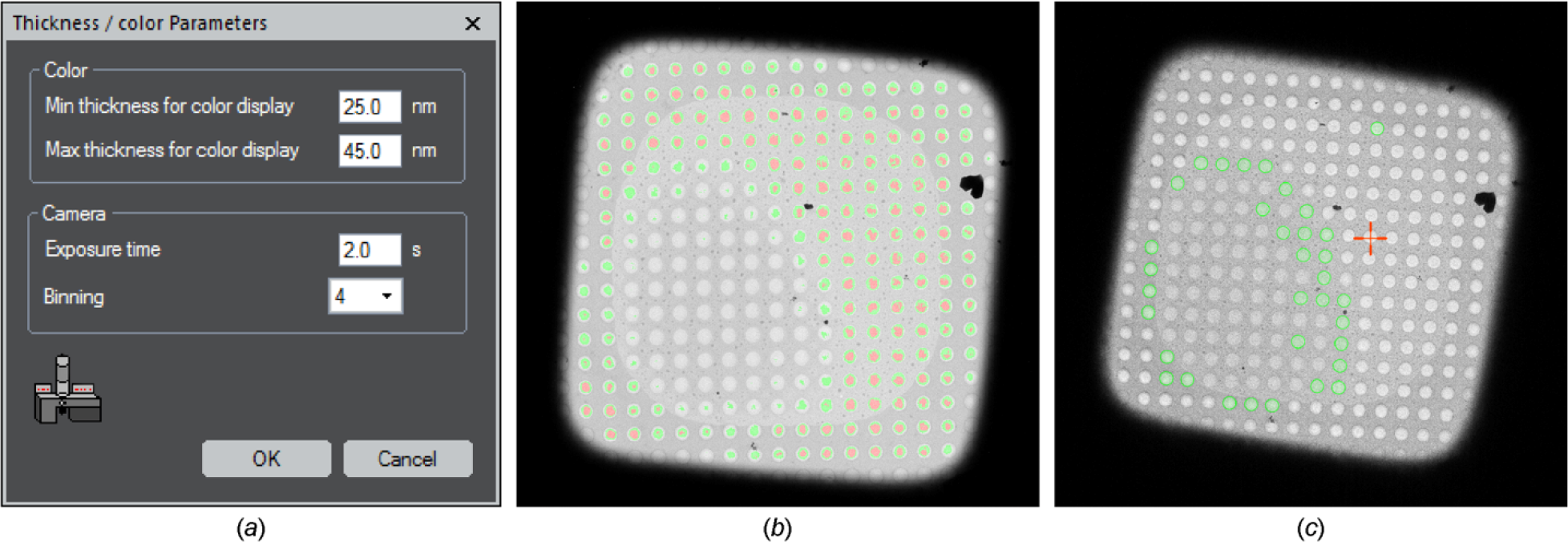
(a) GUI of the Digital Micrograph script, providing access to key parameters, *e.g*., changing the thickness range, even after script execution, (b) Representative color heat map of the sample thickness obtained with the Digital Micrograph script. Coloring shows thickness values above (no color), within (green), and below (red) the user-defined thresholds, (c) Hole targeting of the same square in EPU, where the heat map in (b) was transferred by adjusting the thresholds of the *EPU Filter Ice Quality* histogram.

When executed, the script will collect two images, with and without the energy filter slit inserted. From these two images a ratio image is calculated representing *I/I_zl_* (Eq. 1). In the LM magnification range, where it might be difficult to align the zero-loss peak to the higher magnification, we provided the option of a magnification dependent energy shift to compensate for the offset. The user has to individually determine and provide these shifts in the *Global Tags* (default: 0 eV). For the ALS method, the user needs to define the mean background intensity, which is obtained by measuring the intensity over vacuum within an empty hole (*I_0_*, Eq. 2). This value needs to be determined only once per data collection set-up and is subsequently used to calculate the local thickness (Eq. 2) within a grid square image acquired at the same imaging settings. The switch between the filter-based and ALS method can be activated by a parameter in the *Global Tags*.

The colored heat map is obtained through a pixel-by-pixel calculation (Eq. 1 or Eq.2) and the script assigns each pixel a thickness value. Related to the user defined threshold it will allocate the respective color in an RGB image (too-thin: red, optimal: green, too-thick: no color) (Fig. 1 (b)). The local measurement uses the cursor position in the image and averages the value of the ratio image within a square around these coordinates to calculate the sample thickness. The script also checks the beam intensity to prevent too high coincidence loss (*e.g*., below 10 cnts/px/s for a Gatan K2 camera), thereby ensuring consistent results. While detecting the used voltage, either 200 kV or 300 kV, it will use the respective set of λ_inel_ and λ_ALS_ needed for the calculations. Furthermore, it can distinguish between a K2 and K3 camera (Gatan) and handle the different ways of electron counting. Everything is finished with a GUI that allows to quickly change relevant parameters like the thickness range, as well as exposure time and binning (Fig. 1 (a)). Further parameters can be changed in the *Global Tags* in Digital Micrograph. Since EPU only offers dedicated presets for data collection, and microscope control via Digital Micrograph is limited, we wrote an additional script in JScript within the Thermo Fisher Scientific TEM scripting environment, which allows to store and recall microscope parameters to run the thickness measurement reliably.

Once the sample thickness is determined, the gained knowledge can be transferred into EPU. Here, the *Filter Ice Quality* threshold option during hole targeting is manually adjusted to mimic the hole selection obtained in Digital Micrograph (Fig. 1 (c)). However, this only needs to be done once, for a representative grid square at the beginning when setting up data collection, requiring in total only ~5-10 mins of additional user input. The same parameters will be adopted for any other regions within the same grid and no additional steps are required during data collection. Hence, with the exception that on average fewer holes per grid square are selected and imaged, data acquisition itself is not slowed compared to a conventional setup.

### SerialEM

A clear disadvantage when using EPU as data collection software, is that sample thickness needs to be measured separately in Digital Micrograph first and subsequently transferred into EPU. Moreover, EPU is a commercial software available only for electron microscopes from TFS. To avoid this, we were able to implement the entire workflow in the open-source acquisition software SerialEM (Mastronarde, 2005; Schorb *et al*., 2019) that is compatible with multiple TEM platforms. The newly developed scripts determine the sample thickness and use the acquired values to automatically target holes within the user-defined thickness range. This setup takes advantage of recently added software functions, like *hole finder*, extended by the *hole combiner* for acquisition in multiple holes via beam-image shift. It also requires some newly introduced script commands, which are all available through SerialEM version 3.9 beta1 or higher. Basic parameters (mean free path length, thickness thresholds, imaging settings, correction term, averaging radius) are set as variables in the script to minimize user interaction.

The script will first set the microscope to the desired imaging conditions and acquire an unfiltered and filtered image of a selected grid square. This can be performed for multiple squares and be automatically executed via the *Acquire at Items* function. The newly implemented *hole finder* function then locates the position of each hole within the square. At each hole position the mean intensity in the unfiltered *(*I*)* and filtered image *(I_zl_)* are extracted within a defined radius (Eq. 1), which should be carefully chosen based on the magnification used. This allows to calculate the sample thickness on a hole-to-hole basis that is stored for each item in the *Navigator window* of SerialEM as a *note* (Fig. 2 (a), red box). Based on a user-defined ice thickness range, the calculated values are used to select entries for target acquisition (Fig. 2 (a), green box) and to color positions respectively (too-thick: blue, optimal: green, too-thin: magenta) (Fig. 2 (b)). The acquisition of each grid square requires currently about 5 min, but the process can be automated thanks to the eucentric height routine in SerialEM, such that the user only has to provide the location for the squares of interest. To operate at the LM magnification range, a slightly adapted version of the script is available using the *Search preset* function for image acquisition. This might require an additional energy shift for the respective magnification. As this energy shift is only applied when the slit is inserted and it introduces an image shift, a correction function was implemented allowing to extract the correct intensity values from both the unfiltered and filtered image. For systems without an energy filter we provide a separate script, which can determine the sample thickness using the ALS method outside of a Digital Micrograph implementation. For this the user provides the reference intensity *I_0_* over vacuum within an empty hole, which is divided by the intensity *I* extracted from a grid square image (Eq. 2).

**Figure 2.**
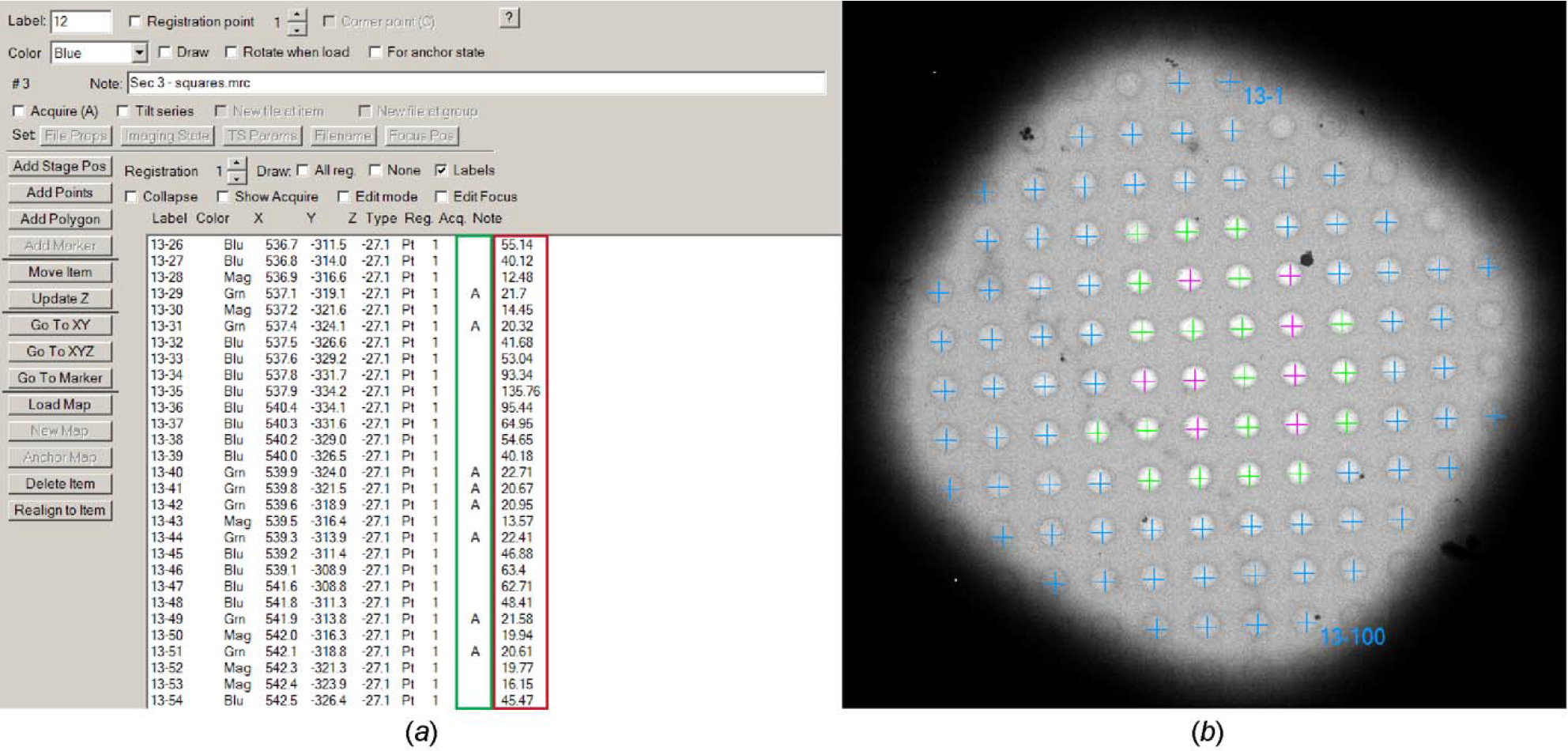
Representative output of the ice thickness script in SerialEM. (a) Navigator window showing the calculated sample thickness in nm for each item (red box) and the selection for target acquisition (green box) based on the predefined thresholds 20 - 40 nm. (b) Hole positions are colored by thickness distinguished into higher (blue), within (green), and lower (magenta) with respect to the thresholds.

The entire workflow is flexible, allowing the user to intervene at any point and make adjustments. If, *e.g*., the initially targeted thickness range is not ideal, a second script allows the user to adjust hole selection using the sample thickness values stored in the *Navigator note* and repeat the threshold-based selection for all entries. This needs minimal user input and finishes within a minute for 5000 points.

### Advantages and limitations

While from our experience - based on small membrane proteins - the optimal sample thickness is in the range of 20 – 40 nm (Fig. 3), this should be tested and optimized for each project, individually. A good starting point is the expected particle size +/− 5-20 nm. For this purpose, on-the-fly pre-processing software packages like FOCUS (Biyani *et al*., 2017), Warp (Tegunov & Cramer, 2018), or Appion (Lander *et al*., 2009) are useful to provide a fast feedback. Here, calculated parameters such as resolution of CTF estimation, particle distribution, or preliminary 2D classifications can be used to assess data quality on the fly and correlate it with the measured sample thickness, allowing the user to define the optimal thickness range and adjust targeting parameters, if required.

**Figure 3.**
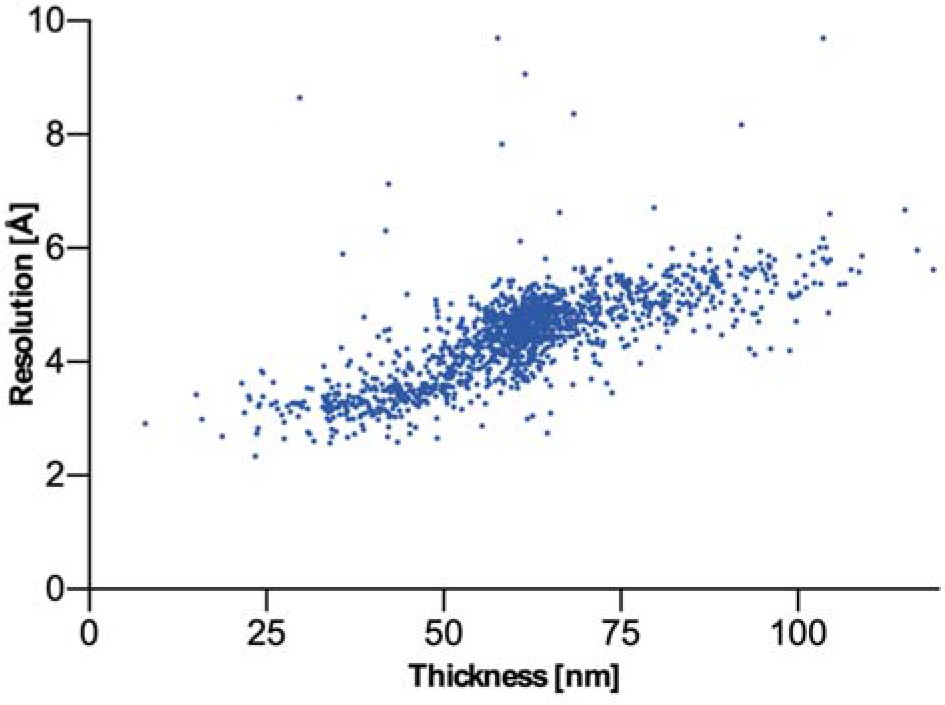
CTF resolution estimation as function of thickness. The graph displays the resolution estimation obtained during CTF determination for over 1400 images with respect to their sample thickness, visualizing the negative effect of thicker areas.

While sample quality and thickness homogeneity can differ significantly between and within grids, the grid squares shown in Figure 1 and 2 are, from our experience, representative. Here, conventional hole selection would have targeted the majority of all visible holes. Yet, more than 50% of the images would not meet the abovementioned quality criteria and be discarded right after data collection or during early image processing. By contrast, our thickness-based approach restricts data collection to only a relatively small fraction of optimal holes present, with 90-95% of the data preserving high-resolution information. The remaining 5-10% of micrographs are usually discarded due to contamination or mismatches between the position of targeting and acquisition. Compared to conventional data acquisition workflows, this approach can, thus, lead to a two- to five-fold increase in retained data.

An emerging technique to speed up data collection, is the use of beam-image shift. It allows to acquire multiple images (*e.g*., 3X3 pattern) by a combined beam and image shift, instead of using stage movement (Cheng *et al*., 2018; Wu *et al*., 2019; Cash *et al*., 2020). The introduced beam tilt can be corrected directly, using the aberration-free image shift (AFIS) correction in EPU (currently only available for TFS Titan Krios) or SerialEM’s own *coma vs. image shift* calibration. Alternatively, it can be corrected later during image processing (Cash *et al*., 2020), by using CTF refinement functions implemented in, *e.g*., RELION or cryoSPARC (Zivanov *et al*., 2018; Punjani *et al*., 2017). Ideally, one would like to combine this approach with the selection of best thickness areas. This is possible in SerialEM with the newly implemented *hole combiner*. It only considers Navigator entries that are selected for acquisition, and groups them in a user defined pattern (*e.g*., 3×3). All entries within a group not marked for acquisition (*i.e*., outside of the defined thickness range) will be skipped. In EPU, with the fast acquisition mode activated, grouping is done automatically, while considering only selected holes within a group. Both setups bring a significant boost in throughput compared to the acquisition by stage movement, while keeping the benefits of optimal area selection.

Another approach aimed to minimize beam-induced movement is the use of gold-coated support films (Russo & Passmore, 2016; Naydenova *et al*., 2020). We can confirm that our workflow also works well with commercially available UltraAuFoil grids (Quantifoil), in particular with the filter-based approach in SerialEM.

The thickness scripts described here, have been developed and tested on TFS microscope systems, namely on a 200kV Talos Arctica and a 300 kV Titan Krios, both equipped with BioQuantum K2/K3. On these systems the energy filter-based method is the preferred option and most accurate in the M or SA magnification range. This works well for a TFS Talos Arctica, as the lowest possible magnification (M910x) still includes a full square of 300 and 400 mesh grids and the majority of a 200 mesh grid square. However, for a TFS Titan Krios, the lowest SA-magnification (SA2250x) covers less than a quarter of a 300 mesh grid square, or about a third of a 400 mesh grid square. Alternatively, a LM-magnification can be selected in combination with the filter based approach. Although slightly less accurate, it provides a bigger field of view able to cover an entire grid square for automatic targeting with SerialEM on a TFS Titan Krios. For systems without an energy filter, the ALS script available for SerialEM, allows to use a flexible magnification range, representing another good alternative.

We noticed, that the measured reference intensity value within an empty carbon film hole was consistently higher than when measured in a fully empty area (*e.g*., broken support film). The nature of this optical effect is not entirely clear to us, but it was more pronounced at 200 kV compared to 300 kV. Since we assume that holes with a vitreous ice layer will also experience this effect, the reference intensity for the ALS method has to be obtained within an empty hole, instead of in an area with no surrounding support film. Furthermore, when measuring empty holes with the filter-based method, the calculated thickness was consistently higher than expected. This led to the introduction of the term C (Eq. 1), which corrects for this offset and brings the measurement back to zero as it should be. At 200 kV this value is around 4 nm for M and SA magnifications, and around 35 nm in the LM range. Notably, this term appears to be unaffected by the defocus setting. For the TFS 300 kV Titan Krios, we estimate a value close to zero (1 nm), both for the M or SA magnification ranges, and about 65 nm for the LM magnification. The values for LM or M/SA can be separately specified and adjusted by the user in the *Global Tags in Digital Micrograph*.

## Summary

The possibility to specifically target only the best areas in a cryo-EM sample, allows the optimization of the data collection workflow. Sample thickness is a key player for cryo-EM data quality, whereby too thick ice lowers the amount of high-resolution information retained, and too thin ice can reduce the number of usable particles in the field of view or affect the structural integrity of the macromolecule of interest. Our open-source scripts allow the user to calculate the thickness in a holey-carbon film, and set up automated data collection targeting only holes within a predefined optimal thickness range. This quality over quantity approach, allows us to restrict imaging to only regions that will provide high-resolution information and thereby avoid the collection of suboptimal images, which would be discarded right after data collection or during early image processing. We were able to successfully implement it in commonly used data acquisition packages and make it compatible with the majority of commercially available TEM’s via the open-source SerialEM.

While only requiring minimal additional user input, this approach maximizes data collection efficiency and lowers the electron microscopy time required per dataset. It is particularly useful, if the speed of data collection is restricted by the microscope hardware and software, or if access time to high-end microscopes, data transfer, data storage and computational power are a bottleneck. For the TFS Talos Arctica, which offers a lower acquisition rate when compared to a TFS Titan Krios (due to higher intrinsic stage drift, which requires longer waiting times, and due to the lack of a third condenser lens, which requires large beam sizes to maintain parallel illumination not allowing multiple data acquisition per hole), this new workflow has demonstrated to be crucial. The approach has been routinely used in all our projects proving its versatility and efficiency experimentally (Garaeva *et al*., 2018; Stock *et al*., 2018; Alvadia *et al*., 2019; Kalienkova *et al*., 2019; Garaeva *et al*., 2019; Arkhipova *et al*., 2020; Sikkema *et al*., 2020; Lam *et al*., 2021).

## Materials and Methods

### Microscopy

The Talos Arctica (Thermo Fisher Scientific) equipped with a BioQuantum/K2 energy filter was operated at 200 kV in zero-loss mode (slit width 20 eV). At M910x magnification (calibrated pixel size 144.6 Å/px) the system was set to microprobe mode with a 50 μm C2 aperture at spot size 8 with gun lens 5 and the C2 lens at 100%, resulting in an electron flux of 12.9 e^−^/px/s (10.7 cnts/px/s). For LM690x magnification (calibrated pixel size 190.7 Å/px) the system was set to microprobe mode with a 50 μm C2 aperture at spot size 7, with gun lens 5 and the C2 lens at 65.0%, resulting in an electron flux of 12.9 e^−^/px/s (10.7 cnts/px/s).

The Titan Krios (Thermo Fisher Scientific) equipped with a BioQuantum/K3 energy filter was operated at 300 kV in zero-loss mode (slit width 20 eV). At SA2250x magnification (calibrated pixel size 37.9 Å/px), the system was set to microprobe mode with a 50 μm C2 aperture at spot size 11, with gun lens 3 and an illuminated area of 40 μm, resulting in an electron flux of 22.6 *e^−^*/px/s. For LM580x magnification (calibrated pixel size 174 Å), the system was set to microprobe mode, with a 50 μm C2 aperture at spot size 8, with gun lens 3 and an illuminated area of 200 μm, resulting in an electron flux of 12.8 e^−^/px/s.

To reduce the noise in the display of the thickness distribution, images are acquired with four-times binning. In Digital Micrograph they are further median filtered to reduce noise. The area of the median filter can be adjusted in the *Global Tags* (default: 3×3 pixel).

### Sample preparation for calibration

Aldolase from rabbit muscle was purified as described previously (Herzik *et al*., 2017). The size exclusion chromatography peak fraction was concentrated to 10 mg/ml, flash frozen in liquid nitrogen and stored at −80 °C. For grid preparation the protein was thawed and first diluted to 2 mg/ml with 20 mM HEPES (pH 7.5), 50 mM NaCl. The diluted solution was further mixed in 1:10 ratio with 10 nm nano gold fiducial suspension resulting in a final protein concentration of 1.8 mg/ml. 2.8 ul of the mixture was applied onto glow discharged (5 mA, 20 sec) R1.2/1.3 holey carbon gold grids (300 mesh, Quantifoil) at 22°C and 100% humidity. The grid was immediately blotted for 2 s and plunge-frozen in liquid ethane/propane using a Vitrobot Mark IV (Thermo Fisher Scientific).

### Calibration

To calibrate λ_inel_, the grids were loaded onto a Talos Arctica operated at 200 kV and equipped with a BioQuantum/K2 (Gatan), slit width 20 eV. A grid square was adjusted to eucentric height and the microscope was set to the imaging conditions for the thickness measurement.

Using a default value for λ_inel_, apparent thickness values for different holes were calculated via the script. A tilt series was acquired at each of the position at SA24000x magnification (calibrated pixel size: 5.5 Å/px) with TOMO (Thermo Fisher Scientific) using a tilt range from - 60° to +60° with 2° increments (dose per tilt angle: 0.6 e^−^/Å^2^, total dose: 36 e^−^/Å^2^). IMOD (Kremer *et al*., 1996; Mastronarde, 1997) was used to reconstruct tomograms using the gold fiducials as marker to align the tilt series. From the cross section of the tomograms the actual ice thickness can be determined. Comparison of the apparent with the actual thickness value at given conditions, allowed to obtain the correct λ_inel_ and λ_ALS_ values, which are used by the Digital Micrograph or the SerialEM script, respectively, when executed for the first time. For the 200 kV Talos Arctica, at a M/SA magnification range, λ_inel (200kV, M/SA)_ is 305 nm. Comparing thickness values at the M magnification with the same grid square at the LM magnification resulted in the respective λ_inel (200kV, LM)_ of 485 nm. For a 300 kV Titan Krios, at the SA magnification range was, the λ_inel (300kV, SA)_ of 435 nm was taken from previous work (Rice *et al*., 2018), which worked very well. The value for the LM range was determine as mentioned above and corresponded to λ_inel (300kV,LM)_ of 805 nm. λ_ALS_ for the LM magnification range was determined by comparison with the filter-based method (600 nm for the TFS 200kV Talos Arctica and 1750 nm for the TFS 300 kV Titan Krios). The values for λ_inel_ and λ_ALS_ seem to be very stable and a recalibration was not necessary at a later point, using the same imaging condition. The fact that we could use reported values for λ_inel_ from Rice et al, 2018 for the Titan Krios, suggests that these values might actually be transferable and stable. However, we do not have sufficient data to ensure a long-term stability and, therefore, recommend to check and eventually recalibrate λ_inel_ and λ_ALS_ after any major change in microscope configuration.

## Acknowledgement

Testing on the Titan Krios was done at the Netherlands Centre for Electron Nanoscopy (NeCEN) with the help of Wen Yang, Rebecca Dillard, and Ludo Renault. We would like to specially thank David N. Mastronarde for providing SerialEM and his dedication to support new functionality, which was crucial to parts of this work, and Michiel Punter for IT support. Special thanks also to Radostin Danev, Craig Yoshioka, Mike Strauss, and Nadav Elad for their helpful discussion on the pre-print version of the manuscript. The work was funded by the NWO Veni grant 722.017.001 and the NWO Start-Up grant 740.018.016 to C.P.

## Data availability

The Digital Micrograph script can be found on the FELMI-ZFE DM-Script Database (https://www.felmi-zfe.at/dm_script/measure-thickness-in-eftem-2/). The script for setting the imaging condition written in JScript is available on GitHub (https://github.com/jrheinberger/SetThicknessMeasurementConditions). All SerialEM scripts have been deposited on the SerialEM Script Repository (https://serialemscripts.nexperion.net/script/63 and related scripts).

Further details regarding installation and execution of the Digital Micrograph script can be found in the supplementary information.

