## Supplementary material for "Optimized cryo-EM data acquisition workflow by sample thickness determination": Manual for Digital Micrograph/EPU workflow (from another version of this preprint: version 2, 2021-03-10)

### Thickness measurement

V 13.0.10

#### Introduction

The DM script can be used to measure ice thickness of a frozen hydrated cryo-TEM specimen. It was developed specifically for single particle data acquisition using EPU (Thermo Fisher Scientific), but it can also be used to estimate local thickness of a specimen for tomography. A colour map can be generated indicating holes with ice of a specified optimal thickness. This allows to set up EPU such that only holes of optimal thickness are selected and used for high-magnification data acquisition.

The standard routine uses the energy filter for thickness measurement, based on the mean free path for inelastic scattering in ice. It works both in M/SA mode and LM mode, as long as the energy shift of the zero-loss peak can be properly corrected, either by microscope alignment or by applying an additional (calibrated) magnification dependent energy shift.

If the filter mode does not work properly in LM, an ALS (Aperture Limited Scattering) mode is available (Rice *et al.*, 2018).

#### Setup of the system

Chose a proper magnification, such that a whole grid square is visible in the image, preferably in the low M (Artica) or low SA range (Krios). Set the illumination (spot size and intensity) such that the number of counts per s and pixel is below 10 for the K2 or 1200 for the K3. This is to avoid too much coincidence loss.

Store the setting in one of the EPU-states, like Zero Loss or Thon Ring, if they are not used by EPU for other purposes. If both are not available you can use the separate Jscript '*SetThicknessMeasurementConditions.hta*' to read, set and store the microscope settings for the thickness measurement.

Make sure that the zero-loss peak (ZLP) remains properly centred when you switch from Acquisition to ThicknessMeasurement state. If necessary, adjust the Cross-over Correction of the microscope (direct alignment) for the low magnification, after centring the ZLP at the higher magnification.

#### Use of the script

- Search for a grid square that is likely to have (too) thin and thicker ice in the holes
- Add the square in EPU > Square Selection; move to the selected square
- Set the eucentric height; again move to the grid square
- In EPU set the 'Zero Loss' or 'Thon Ring' state, depending on how the Thickness Measurement settings were stored (or use the Jscript).

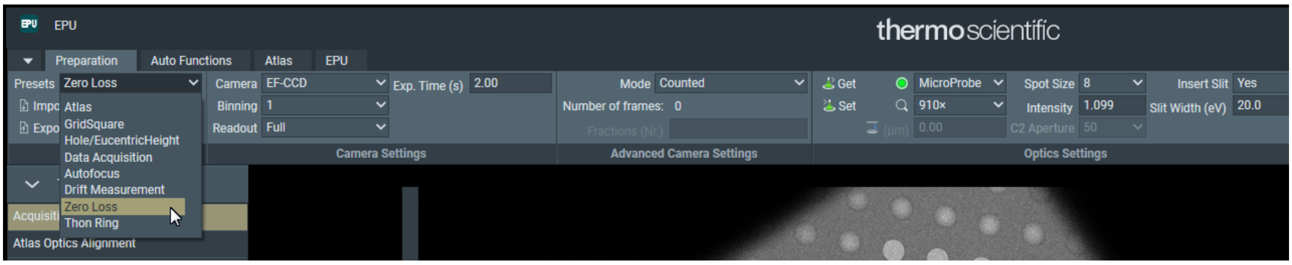

- Choose ‘MeasureThickness [C]’ from the DM > Custom menu

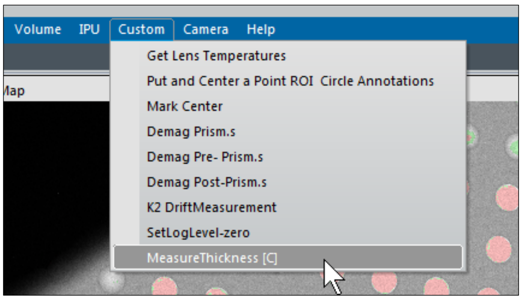

- Two images will be recorded, one with the slit out, one with the slit in. The energy-filtered image is displayed.
- In this image you can do a local thickness measurement by pointing with the mouse and hitting <t> or <T>. The result is printed to the log window.
- You can create a color-coded thickness map by hitting <c> or <C>. The optimal thickness range can be specified at start-up of the routine by holding the <Ctrl> key down, or by hitting <p> (for parameters) when the energy-filtered image or the colour map are displayed.

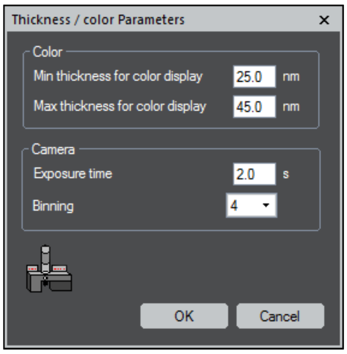

If the colour map is already displayed it will be updated.

- The camera parameters (exposure time and binning) can only be changed at start-up (using the <Ctrl> key).
- The optimal thickness range (between Min and Max) is coloured green, too thin is red, too thick is grey.

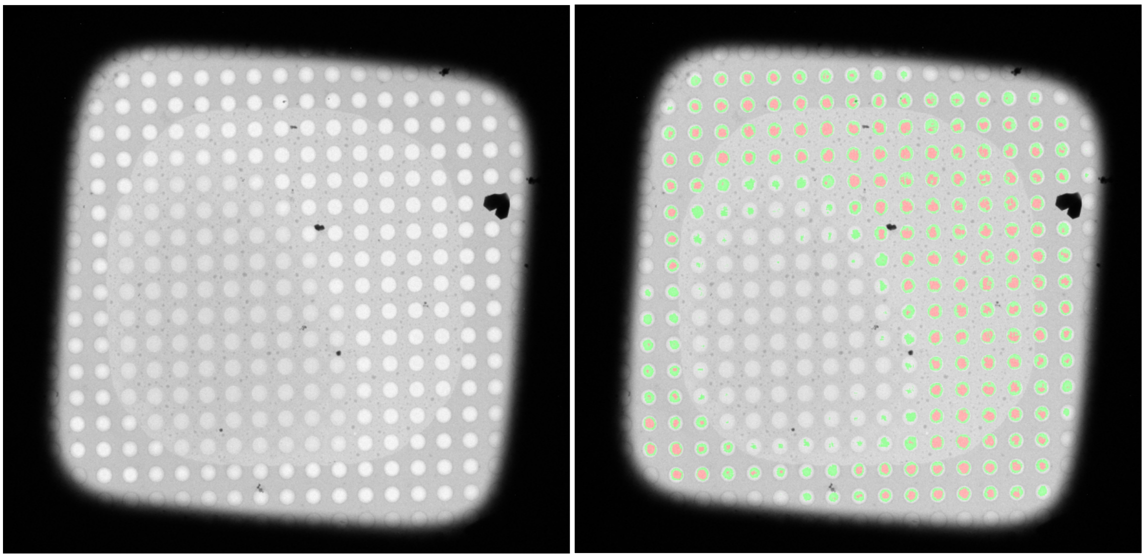

- Note that you can also do local thickness measurements in the colour map like in the energy filtered image using <t>.

- The thickness threshold values can be changed by hitting <p> or <P> (for parameters). If the colour map is already displayed it will be updated.
- Contrast and brightness can be adjusted with the arrow keys: ← and → for brightness, ↑ and ↓ for contrast.
- In EPU set the ‘Grid Square’ state
- In EPU > Hole Selection: Acquire an image
- Measure the Hole Size and Find Holes
- Adjust the Ice Filter such that the selected holes correspond with the holes with the selected thickness from the colour map.

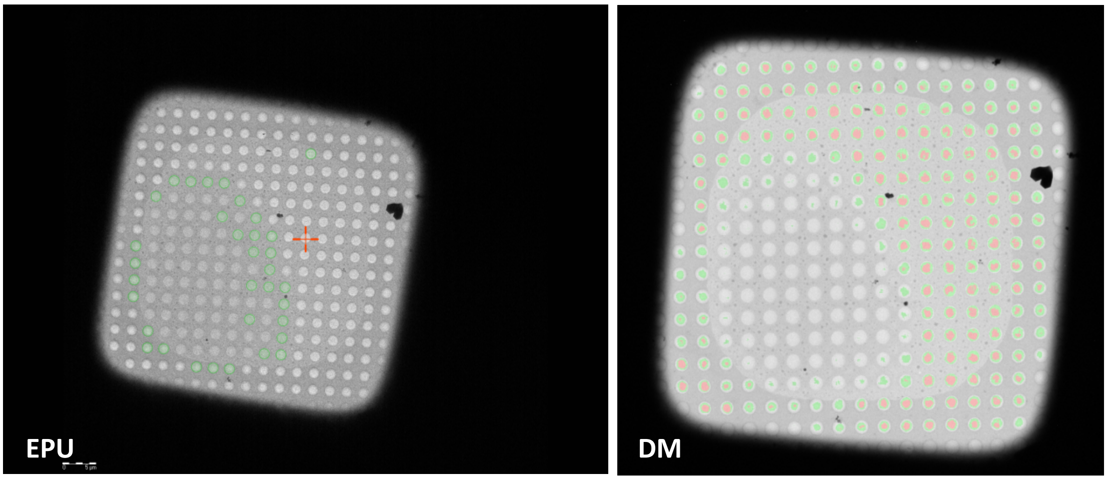

- Always try to include a number (approx. 5) of empty holes in the selection. This gives a reference value of counts/frame to calculate the thickness from the high-magnification images. For 200 kV we use a value of 205 nm for the ALS coefficient (Rice *et al.*, JSB 2018). For 300 kV (and a 100 µm objective aperture) an ALS coefficient of 300 nm is suggested.
- Quit the script on the Gatan computer by hitting the <Space> bar with the pointer outside the image window or hit the <esc> key.

#### Thickness measurement in LM

If the lowest M or SA magnification is too high for the mesh size of the grid used, a magnification in the LM range can be chosen. As mentioned before, check that the ZLP remains centred. If not, try to adjust the cross-over correction. For LM this can only be done in the proper procedure from the ‘Align’ menu. A direct alignment is not available for LM. If it is not possible to centre the ZLP properly by microscope alignment an additional ZeroLossOffset can be defined in a global tag specific for the magnification used:

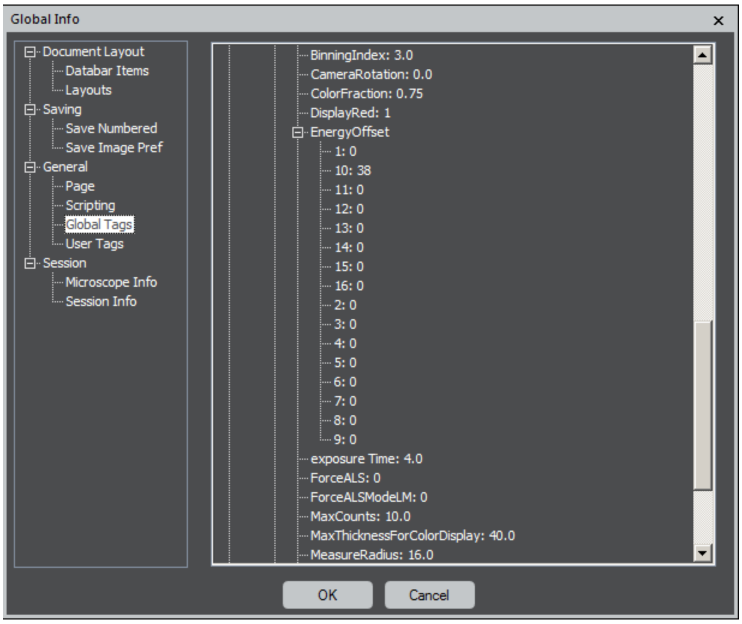

In the example shown a correction of 38 eV is necessary at a magnification of 690x (MagID=10). The table with ZLP correction values is generated (and filled with zeros) the first time the script is run. The current magnification and MagID are printed to the output window for reference.

#### ALS-mode in LM

If the filter mode does not work properly in LM on your system the script can be switched to ALS-mode in LM by setting the global tag 'User:Thickness:EFTEM:ForceALSModeLM' to 1 (the default is zero). The ALS needs a reference intensity in an empty hole to calculate the local thickness. It can be set by pointing the cursor to the centre of an empty hole and hitting <0> on the keyboard. *Note that the intensity measured in an empty hole is generally different from that in a broken mesh of the grid (especially at 200 kV). Setting the reference intensity in a broken mesh or with retracted specimen will lead to unreliable thickness values.*

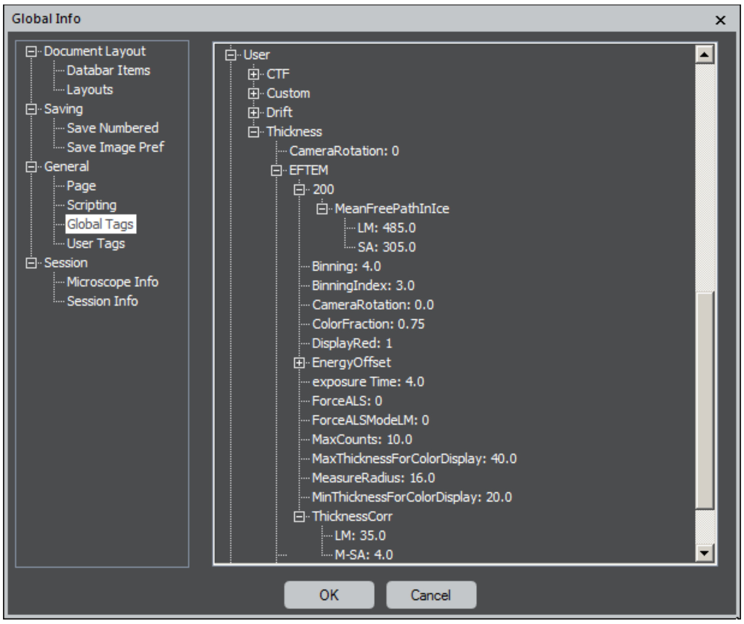

#### Installation

- Open the script in DM. Follow the instructions in the manual to test it. Make sure you have selected a suitable mesh. Click 'Execute' in the script window.
- If the script works properly you can install it in the DM > Custom menu (*File > Install Script*).

The following parameters can be changed as global tags (under "User:Thickness", "User:Thickness:EFTEM" and "User:Thickness:TEM", resp.):

| | |
|---|---|
| CameraRotation | Camera rotation in camera configuration (default 0) |
| MeanFreePathInIce | MFP for 200 or 300 kV and M-SA or LM<br>Default: 200kV M-SA: 305, 200kV LM: 485, 300kV M-SA: 435, 300kV LM: 805 |
| ColorFraction | Strength of the color(s) in the map |
| DisplayRed | Boolean that determines whether too thin areas are marked in red |
| EnergyOffset | For LM. Default 0 |
| ForceALSModeLM | Force ALS mode if filter mode does not work properly in LM |
| MaxCounts | max counts per pixel to avoid too high coincidence loss (default 10 for K2, 1200 for K3 |
| MeasureRadius | Radius of the selection box for local measurements in unbinned pixels |
| ThicknessCorr | Correction in nm to ensure zero thickness in a hole. Default: EFTEM/M-SA 4, EFTEM/LM 35, TEM/M-SA 0, TEM/LM 0 |
| ALSfactor | Default: 200kV M-SA: 800, 200kV LM: 600, 300kV M-SA: 2000, 300kV LM: 1750 |
